# Visualizing Deep Mutational Scan Data

**DOI:** 10.1101/418525

**Authors:** C. K. Sruthi, Hemalatha Balaram, Meher K. Prakash

## Abstract

Site-directed and random mutagenesis are biochemical tools to obtain insights into the structure and function of proteins. Recent advances such as deep mutational scan have allowed a complete scan of all the amino acid positions in a protein with each of the 19 possible alternatives. Mapping out the phenotypic consequences of thousands of single point mutations in the same protein is now possible. Visualizing and analysing the rich data offers an opportunity to learn more about the effects of mutations, for a better understanding and engineering of proteins. This work focuses on such visualization analyses applied to the mutational data of TEM-1 *β*-lactamase. The data is examined in the light of the expected biochemical effects of single point mutations, with the goal of reinforcing or retraining the intuitions. Individual attributes of the amino acid mutations such as the solvent accessible area, charge type change, and distance from the catalytic center capture most of the relevant functional effects. Visualizing the data suggests how combinations of these attributes can be used for a better classification of the effects of mutations, when independently they do not offer a high predictability.

## 1 Introduction

Randomly occurring mutations can cause a loss of structure or/and function of proteins, or provide improved functionality in diverse cellular contexts. The degree of tolerance to mutations at a given position has been used to interpret the role of amino acids in forming or stabilizing the protein structure ^1,2^ or in its function. ^3^Mutational scan evolved as a method to systematically explore the landscape of proteins to understand the sequence-structure-function relationships in proteins, from the perspectives of evolution as well as design. Most mutational scans remained limited to about a hundred mutations or just to a systematic alanine scan. ^4^Alanine being non-bulky and chemically inert was chosen to replace the natively occurring amino acids in 62 positions of human growth hormone to understand its interactions with its receptor. The interpretations which went beyond the structure of interaction complexes, suggested how different amino acids modulated the formation and stability. ^3^

A combination of several factors including a desire to circumvent inherent problems associated with protein purification, development of sequencing technologies, interest in phenotypic screening led to newer methods or rather newer philosophies of mutational scanning. Deep mutational scan ^5^and site saturation mutagenesis ^6^are among the emerging methodologies performing a comprehensive mutation of all the amino acids in a protein to all possible 19 alternatives, and measuring their phenotypic outcomes. ^5^ For example, some recent studies have explored the fitness (dis)advantage in *E. coli* under drug-pressure by performing a few thousands of single-point mutations on *β*-lactamase. ^7^Recent double and triple mutant studies have further pushed the boundaries to the order of hundred thousand independent mutations in a protein. ^8,9^

Deep mutational scan ^5^studies have been generating unprecedented amounts of data, concurrent with all other fields of molecular biology such as next generation sequencing which are undergoing a data explosion as well. The enormous of data from mutagenesis experiments necessitates the development of methods for its analysis, interpretation, and possibly prediction. This data can either be used referentially, seeking the effects of specific mutations or comprehensively to understand the mutational landscape. The focus in the present work is the latter. Mutagenesis experiments in the past decades, with tens or hundreds of mutations on a protein helped develop intuitions on the functional roles of amino acids. With the effects of thousands of mutations now being measured, it will be interesting to see what the new learnings are. Our goal was instead to juxtapose the intuitions from smaller scale experiments over the past decades with those from large scale runs of recent experiments.

Visualizing the data is a way to organize large scale information, note the patterns as well as to observe any deviations from these patterns. Several analyses on the functional interpretation of mutations ^10^and predicting their phenotypic consequences in a diverse range of proteins from ubiquitin to BRCA1,^11^have been developed. Some of the earlier attempts at visualization of deep mutational scan data summarized the entire data in terms of logo-diagrams. ^12^Other approaches combined data from different proteins, summarized and highlighted statistical features such as how likely mutations of amino acid in *α*-helix or *β*-sheet or a turn to be functionally destabilising. ^13,14^ The present work focuses on visualizing deep mutational scan data, with the goal of either reinforcing biochemical intuitions or retraining them by studying the exceptions, possibly contributing towards engineering better proteins. ^15^ We avoided using unsupervised visualization methods used in large scale data analyses, as they may result in abstract relations that may be hard to interpret. Specifically, we focus our visualization analyses on the experimental data ^7,16^ on how the mutations of *β*-lactamase alter the fitness of *E. coli* under different concentrations of ampicilin.

## 2 Methods

The analysis is mainly based on the data obtained from Stiffler *et al.* ^7^, other than in Figure 6 where the deep mutational solubility data from Klesmith *et al.* ^16^ has also been used. The structure of the 263 amino acid long TEM-1 *β*-lactamase was obtained from protein data bank repository (PDB id: 1M40). Hydrogen atoms were added to the structure, using GROMACS ^17^. Solvent accessible surface area (SASA) for each wildtype residue was calculated using *g_sas* tool ^17^ and a probe radius 1.4 Å. SASA was not repeated for the mutations, since it is not easy to predict their stability. Homologous sequences were obtained from Pfam database(PF13354) using PDB id as the query. The sequences obtained were truncated to the reference sequence and the ones with more than 20% gaps were removed, finally having 9105 sequences in the alignment. While studying the effect of a categorical charge type change, amino acids were grouped into four categories - Positively charged (R, H, K), negatively charged (D, E), polar (S, T, N, Q, C) and hydrophobic (A, V, I, L, M, F, Y, W). P and G were not included in any group.

Phenotypic outcome of the mutations used in this analysis was the relative-fitness: ^7^ *F*= *log*_10_(*f ^mutant^/f ^wild−type^*) where *f* is the ratio of allele counts in the selected and unselected population. Zero, negative and positive *F* reflect neutral, loss of function and gain of function mutations respectively. In the analysis where solubility was compared with fitness, we used the deep mutational data from Firnberg *et al.* ^18^. Interestingly, two independent deep scan studies ^7,18^of *β*-lactamase in *E. coli* obtained a non-linear, but highly correlated outcomes (Supplementary Fig. 1) We chose to work with the data of Stiffler *et al.* ^7^as it was 100% complete with all 19 substitutions studied for all wild type amino acids. In several critical scenarios we also validated the conclusions using the other data set. ^18^

## 3 Results

### 3.1 Comprehensive fitness map

A conventional representation of the deep mutational scan data is shown comprehensively and quantitatively in a single palette (Fig. 1A). A quick visual inspection along the two dimensions suggests that a mutation to proline is mostly deleterious regardless of the position or the wildtype amino acid, as well as suggests contiguous clusters of amino acids with high and low sensitivity to substitutions. Also, for a direct comparison of the overall effect of the mutations at each residue position, a panel with sorted fitness effects was used (Supplementary Figure 2). The following intuitive maps were further analysed.

**Fig 1:**
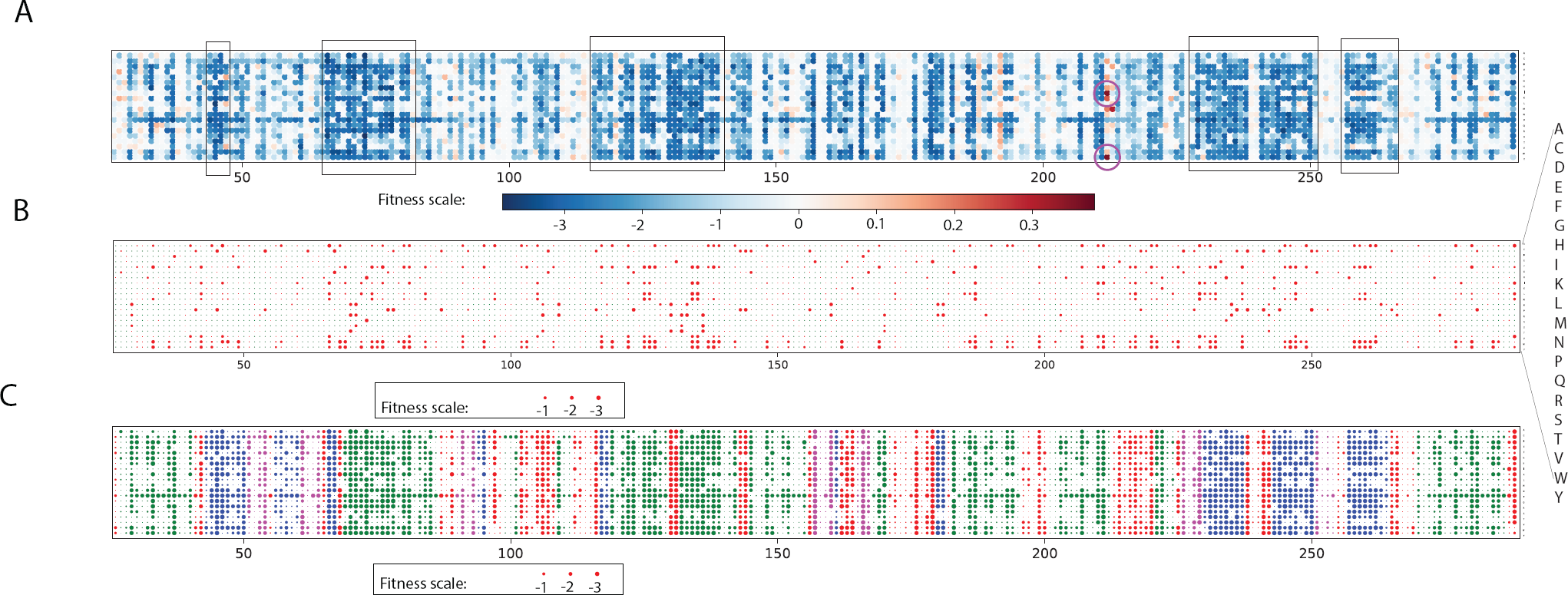
Unexpected changes in fitness maps. **A.** Comprehensive fitness map. Color scale indicates the loss or gain in fitness. Purple circles highlight the only two mutations which show a gain in relative fitness 0.3 (E212I, E212Y). The black boxes indicate continuous regions of the protein that are highly sensitive to mutations. **B.** All the mutations when there is no change in charge type, are shown in red color. It is interesting to note that even in many cases where there is no change in charge type, the fitness is heavily compromised. *Unlike* in Fig. 1A above, the *size* of the data point is used as a measure of the fitness change.**C.** The impact of mutations in different secondary structure: helices (green), *β*-sheets (blue), turn/coil (red), tight turns (magenta) on fitness. The order of amino acids on y-axis is the same for all three subfigures and is shown magnified in **B**.

#### Charge-invariant fitness map

A common intuition is that a substitution involving a charge type change disrupts local interactions or solvent accessibility and leads to a loss of structure and function. Four amino acid categories were considered - positively charged, negatively charged, polar and hydrophobic (**Methods section**). In an attempt to highlight the functional effects that are not intuitively expected, we present the analysis only for the mutations where the charge type of the mutant is the same as the charge type of the wild type, yet the mutation causes a severe loss in fitness (Fig. 1B).

#### Secondary structure fitness map

Data curated from multiple proteins suggested that amino acids in *α*-helices are more tolerant to mutations than those in *β*-sheets. ^14^ In the light of this observation, we examined the sensitivity of - helices, *β*-sheet, turn/coil, tight turns to the deep mutational scan. On average, the map in Fig. 1C reflects that helices are less sensitive to mutations than *β*-sheets (only 55% of the substitutions in helices are deleterious compared to the 80% in *β*-sheets). ^14^Further, as can be expected, loops are least sensitive. It is interesting to note that the contiguous clusters in Fig. 1A have a similarity to the secondary structural patterns.

### 3.2 Functional amino acids

#### Substitution of catalytic residues

Substitution of catalytic residues are expected to be mostly deleterious. Any substitution in the five reported catalytic residues in *β*-lactamase - S70, K73, S130, E166 and A237, other than A237S and A237G leads to high fitness compromise. In catalysis, the backbones of S70 and A237 in conjunction form an oxyanion hole stabilizing a reaction intermediate ^19^, thus tolerating some side chain substitutions at A237. The intolerance of all substitutions except of S and G could probably be because of size constraints.

#### Substitution of binding pocket residues

In addition to the catalytic sites noted above, residues M69, Y105, N132, N170, K234, S235, G236, G238, E240, M272, form the binding pocket. Among these, N132 and K234 are the most sensitive ones as all 19 mutations at these positions result in reduced fitness (*F* < -0.5).

#### Gain of function

Most amino acid mutations cause a loss of function or remain neutral. Only substitutions E212I, E212Y had comparatively high positive relative fitness *F* (*F >* 0.3, *F* is defined in **Methods section**). In both trials ^7^the two mutations have positive effect on fitness, while E212I is reported as beneficial mutation in other studies ^16^ also. It remains to be seen with more careful experimentation if this surprising substitution with charge type change indeed leads to a gain in fitness.

#### Substitution of distal amino acids

From all mutations that lead to a loss of function, ^7^the mutations which also were independently seen to lead to a loss of solubility ^16^ were eliminated. 57 substitutions of 16 wildtype amino acids were more than 15 Å away from the catalytic residues and yet lead to *F* < −1.5.

### 3.3 Comparison with structure and sequence data

#### Conservation versus fitness

Typically, evolutionary conservation reflects the functional importance of an amino acid. Fig. 2A shows the relation between conservation and the fitness effects from deep scan data of TEM-1 *β*-lactamase (alanine scan data is shown in Supplementary Fig. 3). Conservation alone does not resolve the fitness effect completely. However, as suggested by the guideline in Fig. 2A, there is a reduction in fitness when conserved amino acids are mutated. We highlight two types of exceptions to the expected intuitions: (1) The amino acids that have less than 20% conservation and yet severely affect function upon mutation (F < -1). The mutations N52C, K55(C,P), E58(C,F,H,I,L,M,P,V,W,Y), S82(C, P), S98P, N100C, T140P, T141(F,K,P,W,Y), E197(F, L), P219(F,I,W,Y), F230(C,D,E,G,I,K,L,N,P,Q,R,S,T) and S258P are deleterious even though the wild type residue is poorly conserved. All these substitutions are away from the catalytic sites and other than S82C,F230I and F230L, involved a charge type change. Interestingly, most of these substitutions also involved a loss of solubility ^16^ which could be the reason for reduced functional fitness. (2) The amino acids were conserved (*>* 80%) but their substitution did not affect the function significantly. G156D, G156E, G156N and G236A are the substitutions which are neutral despite high conservation, and the charge type of the mutant differs from the wild type. As conservation quantifies only variability at a specific position and does not distinguish different substitutions, we calculated position specific scoring matrix(PSSM) using PSI-BLAST and explored its relation with fitness. We observe only a weak correlation (Supplementary Fig. 5).

**Fig 2:**
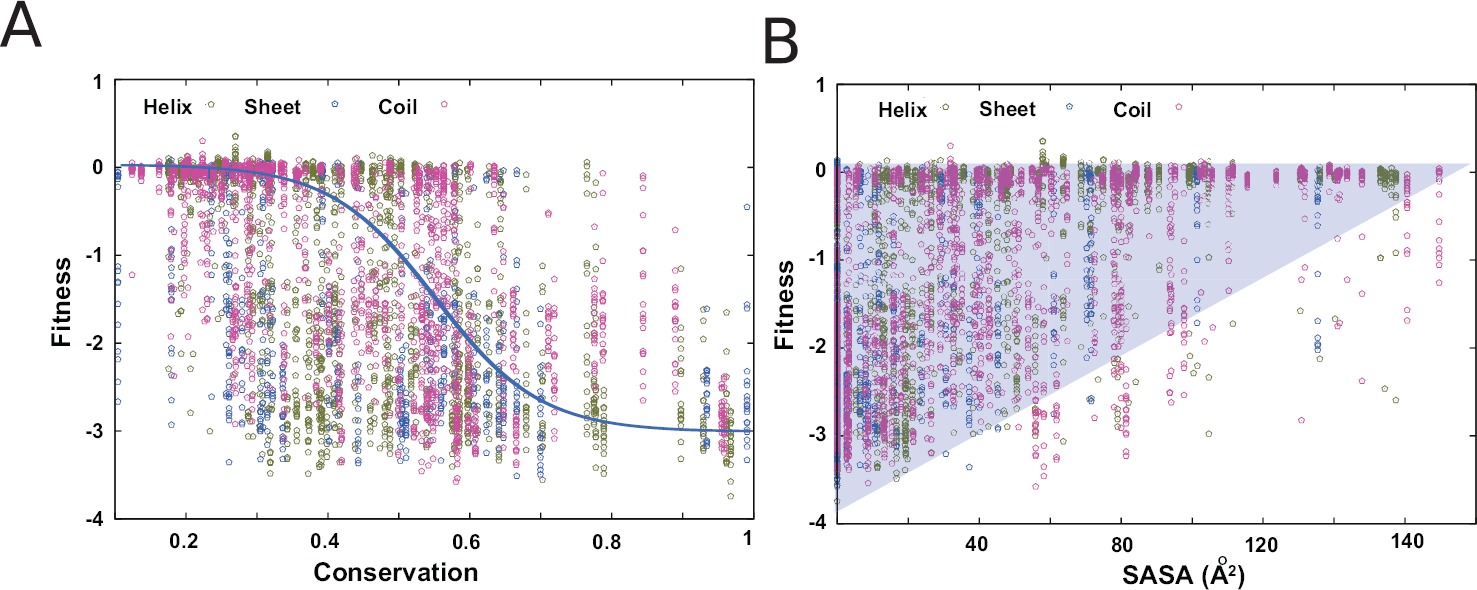
Structure and Sequence Comparison. **A.** The relationship between conservation and fitness was studied using the homologous sequences for TEM-1 *β*-lactamase (Pfam id PF13354). It can be seen that the number of neutral substitutions decrease considerably for amino acids with conservation > 60%. The blue guideline is drawn to highlight the sigmoidal pattern. **B.** SASA vs. fitness showed an interesting half-triangular pattern. The trend is even strongly observed in the alanine-scan mutation (Supplementary Figure 3). The deviations from this half-triangular pattern are noted in the main text.

#### Solvent accessible surface area (SASA) versus fitness

The relation between fitness and SASA of wild-type amino acid which reflects how buried the amino acids are is shown in Fig. 2B (alanine scan results in Supplementary Fig. 3) The intuitive learning from this figure is that substitutions at amino acids which are completely buried can potentially range from neutral to deleterious, while the effect tapers off for amino acids with high SASA values which have minimal effect on fitness. The mutations defining the frontier and showing the highest fitness compromise at any given SASA, were recorded by taking note of the alanine scan mutations near the triangular border in the plot. Of the amino acids P27, L57, R61, R65, F66, S70, K73, R93, Y105, S130, N132, N136, D157, R161, E166, R222, W229 and W290, which are on the frontier of highest fitness loss, most are near the binding pocket. W229 has been known to act allosterically ^20^. However, the reasons for the functional compromise of mutations at P27 and R222 are not clear to us.

### 3.4 Representative scans

#### non-ALA scan

Alanine scan evolved as a popular biochemical tool, representative of all other substitutions. Where alanine scan suggests that the mutations are strongly deleterious (*F* < −2), the fitness effects of all other amino acid substitutions were aligned with it. On the contrary, we also noted the substitutions of the amino acids C77, T109, G236, G238, G245, G283 which are deleterious when the corresponding alanine scan suggests them to be neutral (Fig. 3A). Thus, while alanine scan has been adopted as a standard for the practical reasons of simplicity, it seems to be representative of the effect of mutations in general.

**Fig 3:**
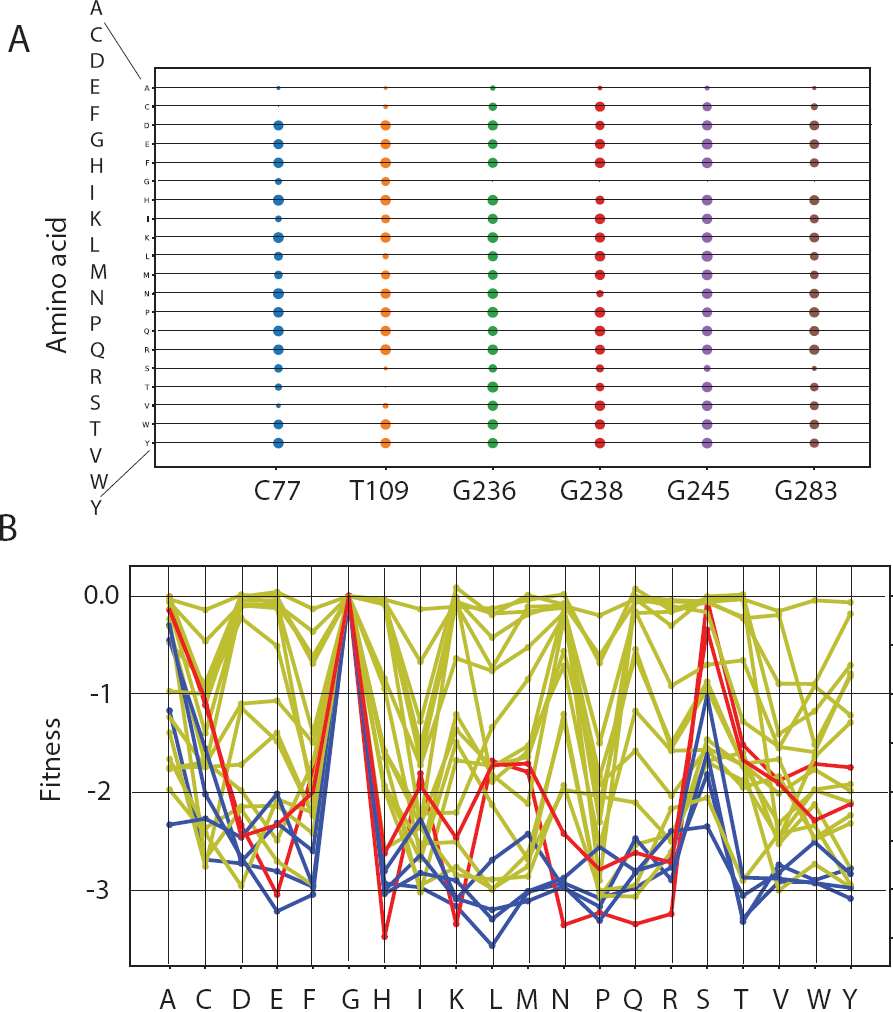
Representative scans. **A.** The data points represent the positions where the effect of alanine-scan was found to be opposite to the replacements with all other amino acids. The size of the data points represents the intensity of fitness loss. **B.** This graph represents the change in fitness when the glycines in the wild type are replaced by all 19 alternatives. Color represents the secondary structure (red-helix, blue-*β*-sheet, yellow-turn/coil)

#### Replacement of glycine

Since in Fig. 3A, the mutations of naturally occurring glycine seems to be counter-intuitive relative to alanine scan, and glycine is specifically known to be a helix breaker, we further analysed the role of secondary structure on the replacement of glycine in Fig. 3B. On average, it appears that when glycine is not in helix or *β*-sheet, the residue is less sensitive to mutations.

#### Substitutions to and from proline

Proline being an imino acid can not donate hydrogen, and it usually plays an important role as a secondary structure breaker. It can be seen from Fig. 1A that most of the substitutions to proline are deleterious (225 out of 251 positions where the wild type is not proline). For 150 of these residues, which are in either helices or sheets, this effect may be expected due to the loss of hydrogen bonds. Of the remaining 75 that are in coil or turn, 61 have *Φ* angle differing from the -60° assumed by prolines by more than 10°. The neutral substitutions were mostly in turns, coils or at the ends of the secondary structures (other than N100, K146, E147, A202, D273). Proline is the only amino acid which can assume *cis* conformation and help forming turns in amino acid chains. Hence mutations of *cis*-P can potentially affect the ability to form structures. We analysed the fitness effect of the substitution of the only *cis*-P167 and surprisingly found not to have very crucial role in structure as it tolerates mutations to I, L, T and V.

### 3.5 Spatial organization of mutational sensitivity

#### Highly sensitive amino acids

The amino acids which could not be replaced at all by any of the other amino acids without a strong fitness compromise (F < -1.5): F66, S70, T71, K73, S130, D131, N132, N136, D157, E166, D179, K234, G242, G251 were noted (Supplementary Fig. 6). Most of these highly sensitive amino acids are either the catalytic residues (S70, K73, S130 and E166) or near to the catalytic residues (distance from any one of the catalytic residues < 10 Å). Exceptions to this observation are F66 (12.7 Å), D157 (17.2 Å) and G251 (22.5 Å). Comparing with the solubility data, ^16^ it can be seen that mutations at F66 in general affects the solubility (or folding) and hence the function. It has been proposed that volume constraints could be one of the reasons for the deleterious effects of substitutions at this position ^21^. D157 is involved in internal electrostatic interactions stabilizing the loop between the first and second *β* sheets ^22^. The solubility data also supports the role of D157 in maintaining the structural stability. Residue position G251 is interesting as it is far from from the catalytic site and also not all mutations at this site affect stability. To our knowledge, no studies have clarified the role of this residue in protein function.

#### Least sensitive amino acids

We also examined the residues which can tolerate all 19 substitutions by allowing for a loss of fitness up to *F ≥ −*0.5. As expected, all these resilient positions: E28, T114, K146, E147, H158, P174, E177, L198, L201, A227, E240, D254, Q269, D273, shown in Supplementary Fig. 6 are on the surface of the protein.

#### Clustering of positions

The comprehensive fitness map with 20 possible amino acids on y-axis and all 263 amino acids on x-axis were clustered based on fitness profiles. Interestingly the y-axis clusters show that proline is completely different from all other amino acids, aromatic amino acids cluster together. The two major clusters on x-axis, separate the amino acids into buried and exposed amino acids, reiterating the high and low sensitivity groups noted above. The clustering of the data thus naturally separates the buried and solvent exposed amino acids. One major cluster with 62 residues includes the catalytic and most of the the binding pocket residues.

### 3.6 Role of Packing

Δ**Volume and SASA:** The change in volume upon mutation of amino acids using their standard volumes ^23^, at different levels of solvent accessibility, was explored (Fig. 5). The data on average supports the intuition that amino acids which are completely buried and have a zero or reduced solvent accessible area do not tolerate mutations. At intermediate solvent accessibility conditions, interestingly, a reduction in volume of the amino acid seems to be more deleterious in general. This could be because of the cavities being created which affects the packing of the residues. It is known that cavity creating mutations reduce the stability of proteins. ^24^ The number of atom-atom contacts (with 4 Å cut-off) was also plotted against the fitness (Supplementary Fig. 7).

**Fig 4:**
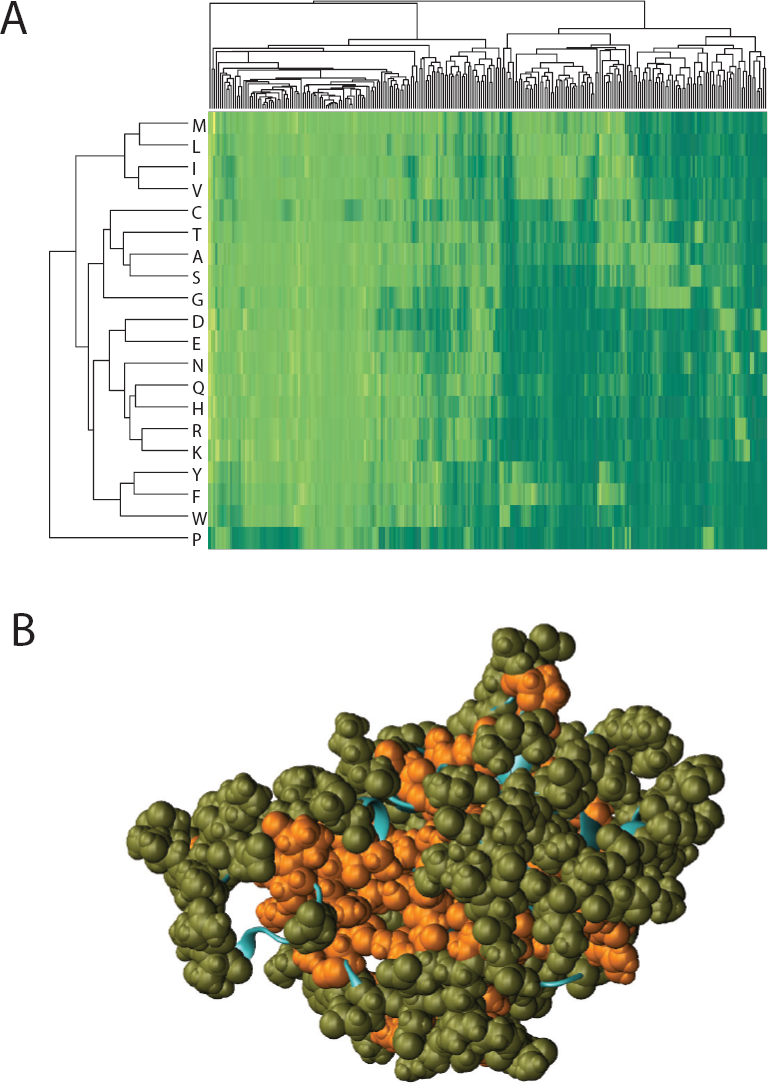
Spatial organization of mutational sensitivity. **A.** Clustering the comprehensive fitness data shown in Fig. 1A using the clust routine of Matlab created simultaneous hierarchies both among the 20 amino acid types, as well as among the 263 different amino acid positions. Substitutions to different amino acids can be intuitively grouped as those to proline, and aromatic amino acids, etc **B.** Of the four major clusters of amino acid positions, the most sensitive (orange) and the least sensitive (green) are shown, which are buried from and exposed to solvent respectively.

**Fig 5:**
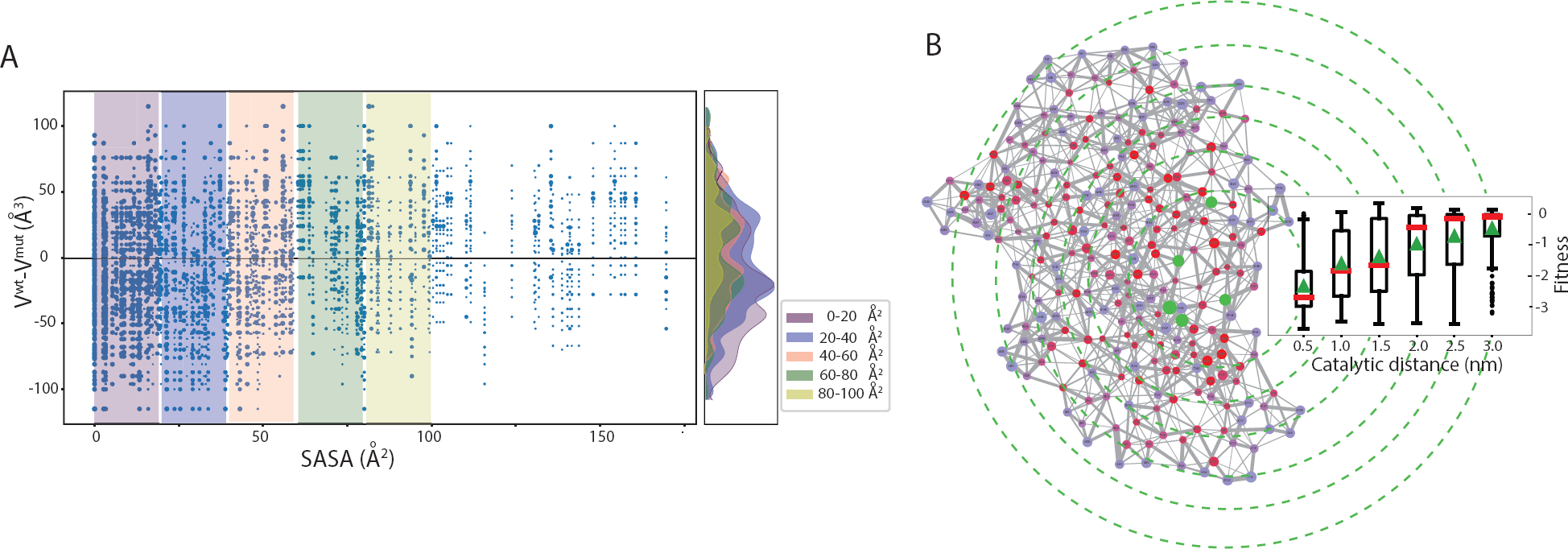
Role of packing. **A.** The fitness compromise (size of the points) is shown relative to the solvent accessible surface area of the wild-type amino acid and the change in volume upon mutation. The histograms on the right show that the distribution of mutations shifts from increase of volume mutations at smaller SASA to cavity creating mutations at intermediate SASA values. **B.** The average fitness effect of substitution on each residue is shown as color (red being the strongest and blue the weakest) on the amino acid contact network. Catalyitc residues are highlighted in green color. Nodes correspond to amino acids. The thickness of edges is proportional to the number of contacts and node size is inversely related to the standard deviation of the fitness for all 19 substitutions. Box plots for the distribution of fitness values for mutations of residues at different distances from the catalytic residues, (schematically represented by the circles) is shown. The green triangle and red line correspond to the mean and median of the fitness respectively.

While the average trends are intuitive, like substitutions of residues with higher number of contacts result in larger fitness effect, the variation in fitness for a given number of contacts are high. Further, the average fitness compromise appears to be directly related to the distance from the catalytic site (Fig. 5B). We also examined if any of the amino acids in the disallowed region of the Ramachandran map as defined in Gunasekaran *et al.* ^25^ had significantly higher impact on the fitness (Supplementary Figure 4). The residues identified outside the classical allowed region were M69, H29 and L220. All substitutions of amino acids L220 in the disallowed region affect the functional fitness significantly. From the solubility data L220 does not seem to contribute in maintaining the structure.

### 3.7 Structural stability and solubility

#### Thermodynamic stability

Using the prevalence of the different amino acids in *α*-helix and *β*-sheet, as a surrogate for thermodynamic stability (Δ*G*), we estimated the stability consequence of a mutation (ΔΔG) for the amino acids in these secondary structures ^26,27^. ΔΔ*G* was uncorrelated with the loss of function (Supplementary Fig. 8), possibly for several reasons including the lack of a direct relation between prevalence and thermodynamic stability or because thermodynamic stability may not define the functional fitness at physiological conditions. We made another comparison of the functional fitness with the solubility data from other experiments ^16^ and distance from the catalytic center (Fig. 6) hoping for a better correlation. The trend we see is logical apart from the very few mutations which appear in the region where solubility < 0 and fitness 1 which corresponds to misfolded structure and improved function. The charge type changing substitutions shown as larger points but do not show any difference relative to no-charge type change (smaller points).

**Fig 6:**
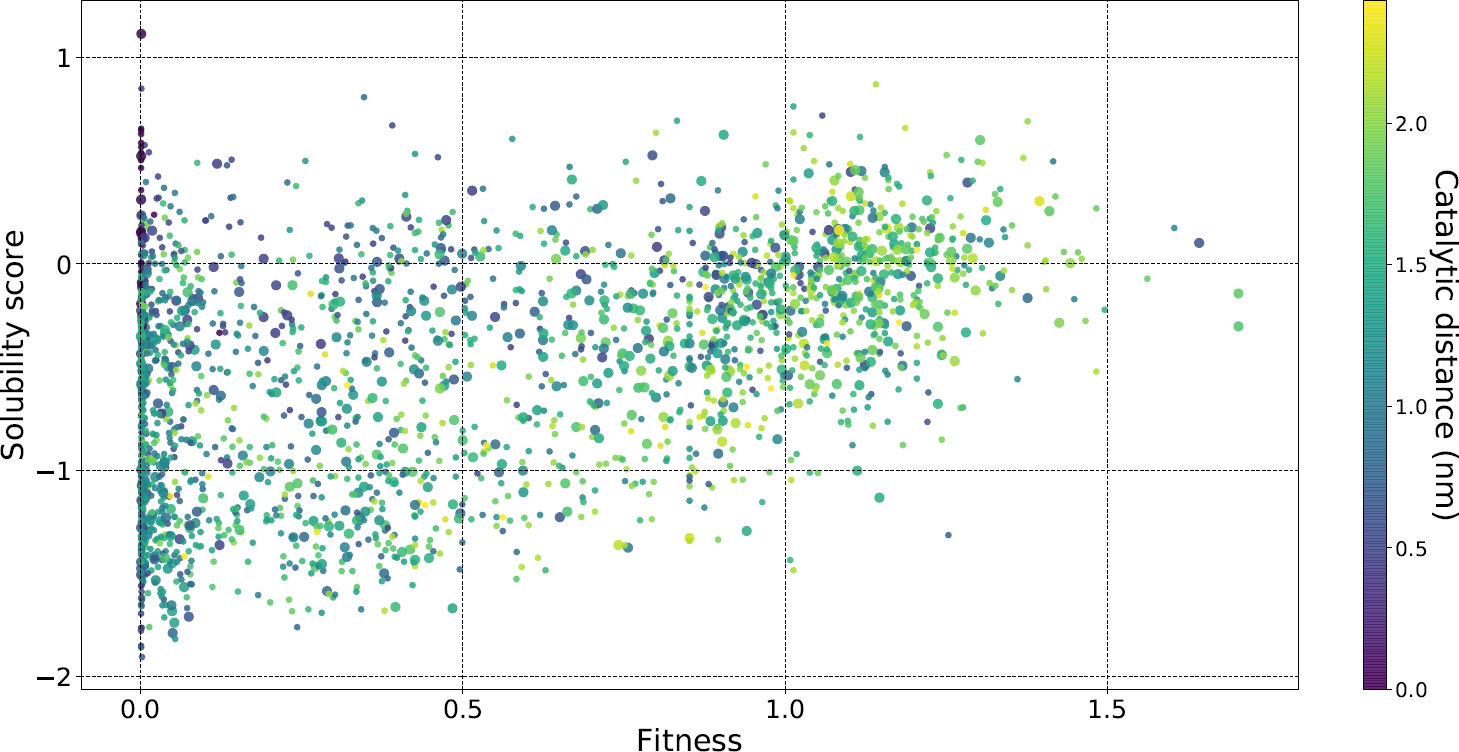
Role of solubity. Deep mutational solubility data from Klesmith *et al.* ^16^ and deep mutational fitness data from Firnberg et al. ^18^ are plotted against each other. Negative, zero and positive values on the solubility scale correspond to reduced, same and increased solubility respectively compared to the wild type. In the scale of Firnberg *et al.*, ^18^ fitness 1 is neutral and more and less than this suggest beneficial and deleterious mutations respectively. The color indicates the distance from the catalytic center(nm). Two data point sizes are used. Smaller size is when the mutation did not change the charge type and the larger size when the mutation changed it. No pattern was observed with respect to the change in the charge type after substitution.

#### Disulfide bonds

The disulfide bond C77-C123 is highly conserved among *β*-lactamases and it is believed to be structurally important. All mutations of C123 were deleterious where as C77(A,V) are tolerated. The slight difference in the sensitivity between the substitutions at C123 and C77, was consistent across the two studies ^7,18^. The solubility of the two mutants is not compromised either, ^16^ suggesting that this disulfide bond is not critical for structural stability. The tolerance of two substitutions, both of which are hydrophobic, suggests that probably the local environment of the protein does not tolerate the lone charge appearing on C77, when C123 is substituted.

#### Salt bridges

Functional importance of amino acids involved in salt bridges showed no correlation with formation of salt bridge (Supplementary Fig. 9). We further compared the sensitivities of the four lysines which formed salt bridges to the others which did not (Supplementary Fig. 10). We find that the most deleterious substitutions depended on the distance from the catalytic site, rather than on the salt bridges.

#### Aromatic residues

To identify crucial aromatic interactions, we analysed the fitness effects of residues F, Y and W. The positions W210, W229, W290 and F66 do not tolerate any mutations. W229 and W290 are involved in a T-shaped pi-stacking, and this sensitivity is expected based on the structural intuitions.

### 3.8 Dose-response curve

Deep mutational scan experiment data from six ampicillin concentrations was combined to generate dose-response curves for the different mutations. Supplementary Fig. 11 clusters each of the thousands of possible substitutions, and their corresponding phenotypic response at six drug concentrations in the *β*-lactamase. One observation from these dose-response curves is the continuity of the data, across the increasing concentrations, which allows extrapolating the experiments or calculations from one concentration, say 2500 *µ*g/ml that is several orders of magnitude higher than the physiologically relevant minimum inhibitory concentration of about 0.0625 *µ*g/ml ^28^, to lower concentrations which are more relevant physiologically but with noisy experimental data. Also the clusters of different levels of sensitivity identified at high concentrations in Fig. 7 continue to be relevant at lower concentrations as well.

**Fig 7:**
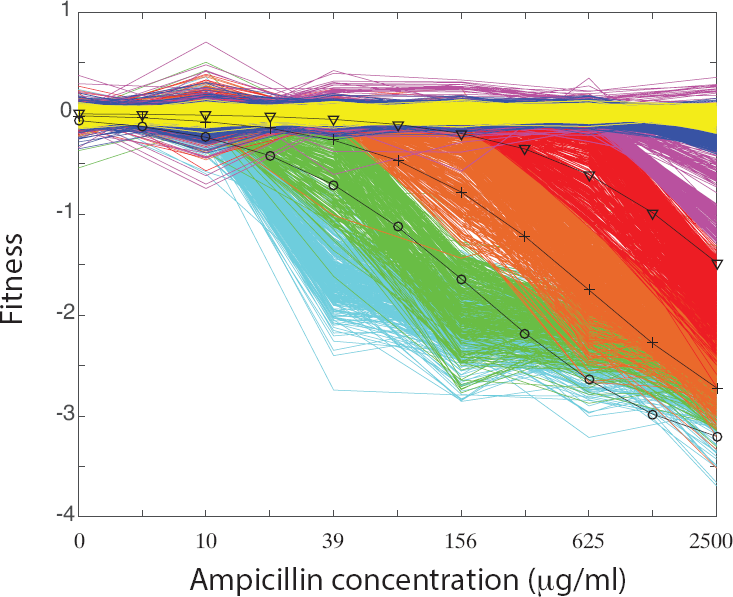
Dose-response curve. The data obtained from multiple concentrations was clustered. Along the x-axis are the six experimentally measured concentrations and y-axis are the 5000 mutations on which measurements were made. The clustered data was used to create bundles of mutations with different levels of sensitivity to the drug concentration. The different dotted lines which are sigmoidal functions suggest a monotonous and predictable dose-response.

## 4 Discussion

The visualization analyses in this work had two goals: Firstly to explore different ways of presenting the results from about 5000 mutations, each studied at 6 different drug concentrations. Secondly, to summarize the observations in a way to juxtapose with the biochemical intuitions. The phenotypic effect in a *in-cellulo* context is a downstream effect of the mutations which perturb the structural stability, and interactions with the substrates or other proteins. Despite this cascading effect, interestingly, the consequences of most of the mutations could be classified using biochemical intuitions.

The fitness loss depended on the conservation or the SASA of the wild type amino acid with characteristic sigmoidal or triangular patterns. The contiguous clusters of amino acids with similar sensitivities on average follows the secondary structural patterns. The dose-sensitivity of the mutants follows monotonous sigmoidal behavior over a few orders of magnitude difference in dosage quantities, opening the possibility of performing the experiments at higher drug concentrations and inferring them for physiologically relevant other concentrations. Functionally relevant amino acids seem mostly buried, and clustered around the catalytic site, even up to 15Å around it, and alanine scan almost faithfully reflected the consequences of most other amino acid substitutions. Interestingly the only few mutations that may be interpreted as gain of function candidate mutations, after correcting for the experimental errors involved a change in charge-type, and were further away from the catalytic site. While *β*-lactamase is believed to be a monomer in the functional context, we examined potential dimer interfaces ^29^, and they did not involve E212. Thus the role of E212 in gain of function still open.

A protein is complicated, and predicting its structure or function is non-trivial. Clearly a single biophysical or biochemical parameter can not capture the consequence of a mutation in its entirety. While most of the biochemical intuitions were obeyed by the thousands of mutations, clearly any of the graphics showed several exceptions. In each individual graphic presented in the visualization analyses, we also explored the possible meaning of the exception by further analysing them for other biochemical intuitions. Thus it is clear that a combination of several biochemical intuitions does help the classification of the functional effect (Supplementary Fig. 12). The same was explored more systematically in Fig. 8 in an analysis on the subset (2892 mutations) of the experimental data where the substitutions had a fitness compromise (*F* < −0.5). Each mutation was independently classified as neutral or deleterious using different biochemical parameters and their corresponding threshold values. The numbers appearing in the different overlap regions in Fig. 8 indicate the number of mutations for which the variables defining the overlap all result in a false-neutral prediction. The interesting region in the center shows that when all 5 variables classify a mutation as neutral there were no false predictions. If only the three structural variables corresponding to the distance from the catalytic site, number of contacts and SASA were used together, the number of false-neutral predictions are 66, thus requiring the sequence related information also to reach a better classification.

**Fig. 8.**
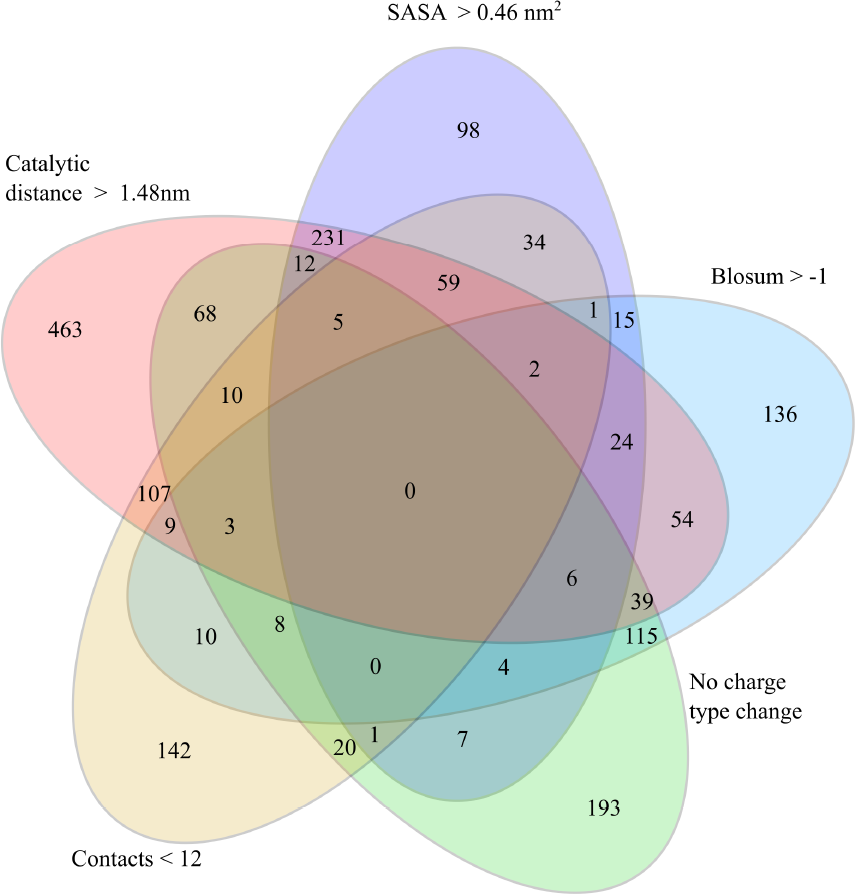

Venn diagram showing the number of substitutions that do not follow the intuitions related to the different structural and sequence related biochemical characteristics of the wild type and the substituting amino acids. Different thresholds indicated in the figure were used on each of the individual parameters, to classify the mutations as being deleterious or neutral. The central zero indicates no false-neutral predictions when all five variables classify the mutation as neutral.

## 5 Conclusions

Different visual representations were used to visualize, analyse and summarize the large scale mutational data on the same protein. One of the core goals of the visualization analyses was to make a direct comparison with the biochemical intuitions gained from several small scale experiments over the past decades. It appeared that some of the results were counter intuitive, however, by using a combination of the several intuitions, the possible deleterious effects of the mutations could be classified. Visualization thus presents a way to summarize, analyse and classify the effect of mutations. Further visualizations with the effects on solubility, and gain-of-fitness mutations, when more such data with smaller variance becomes available, will lead to enhance the intuitions about the interplay of protein stability and function.

## 6 Acknowledgments

We thank Prof. Tim Whitehead and Dr. Justin Klesmith for clarifications regarding their deep scan solubility studies on *β*-lactamase.

## Supplementary Information

### Data Used

Deep mutational scan studies on *β* lactamase were independently performed by two different groups, Stiffler *et al.* ^7^ and Firnberg *et al.* ^18^ We compared the two data sets for consistency across the thousands of mutations, in Supplementary Fig. 1. The data from both the sets was well correlated, although in a very non-linear way. The data set of Stiffler *et al.* ^7^ was 100% complete, in the sense that for every amino acid residue that has been mutated, data on all substitutions was available. So we used this data for the present analysis.

**Supplementary Fig. 1:**
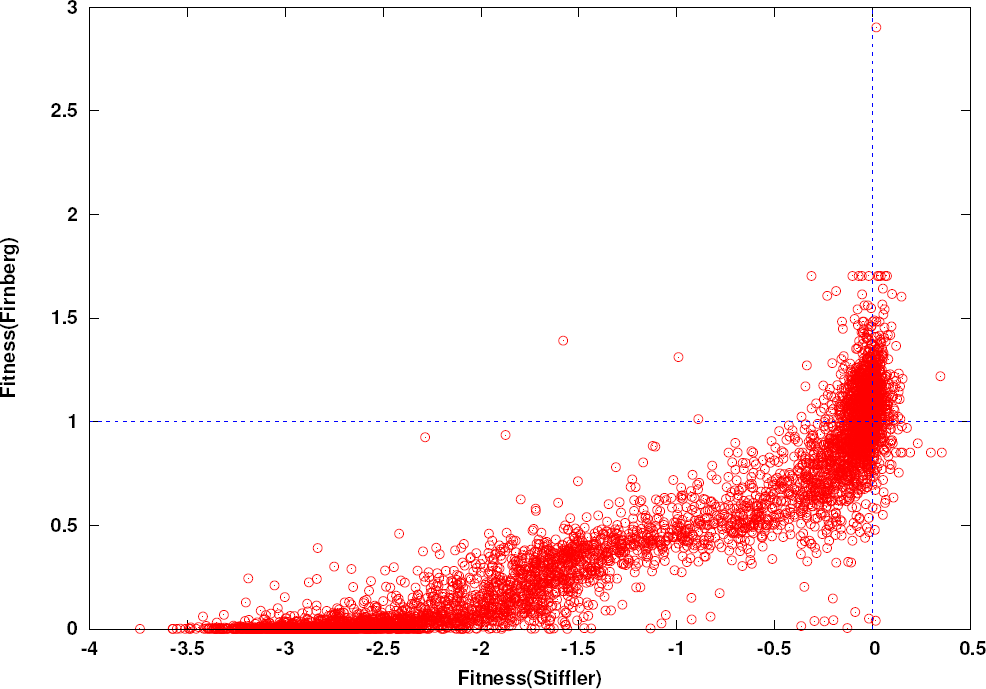
A comparison of the results reported in two different deep mutational scan experiments where the mutations in beta-lactamase and their fitness effects on *E. coli* were studied. The two dashed lines passing through 0 and 1 represent the wildtype fitness in the Stiffler *et al.* ^7^ and Firnberg *et al.* ^18^ experiments respectively.

### Overall intensity of substitutions

In the comprehensive fitness panels shown in Fig. 1 of the main article, the data is presented in a fairly linear fashion, with the amino number along one direction and the 20 amino acid substitutions sorted in an alphabetical order. There can also be other ways to organize the substitutions, say by grouping all amino acid substitutions of a charge type together, which allows easy comparisons of substitutions to the same charge type. Here in Supplementary Fig. 2, we show another representation which is meant mainly to compare the overall intensity of fitness compromise with the 19 amino acid substitutions. The 20 data points corresponding to each residue position in the protein was sorted in an increasing order of loss of fitness. The graphic representation makes it easy to identify residues which have high, medium and low sensitivities to mutations. We see that there are several contiguous segments where all the amino acids are extremely sensitive to mutations.

**Supplementary Fig. 2:**
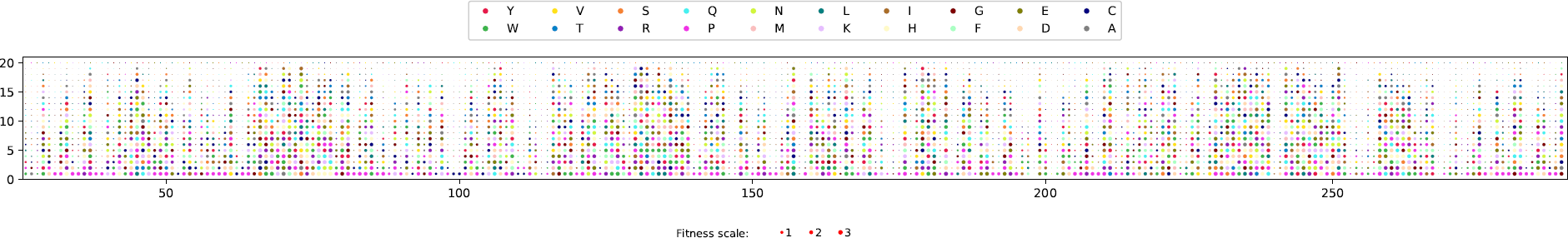
Fitness effects of substitutions at each position is sorted and is shown in the colormap. The color indicates the substitute amino acid and the size indicates the magnitude of fitness effect.

### Relation with conservation and solvent exposure

Conservation and the extent by which an amino acid is buried are commonly associated with functional role. These intuitions are based on a handful of mutations. We examine these hypotheses, with the goal of quantifying them, in the light of the large scale deep mutational data. The complete data is shown in Fig. 2 of the main article. Since it was seen that alanine scan is representative of the complete mutational scan, for the sake of simpler interpretations, here we examined the quantitative relations only with the alanine scan. The analysis reflects sigmoidal and triangular patterns with conservation and solvent exposure.

**Supplementary Fig. 3:**
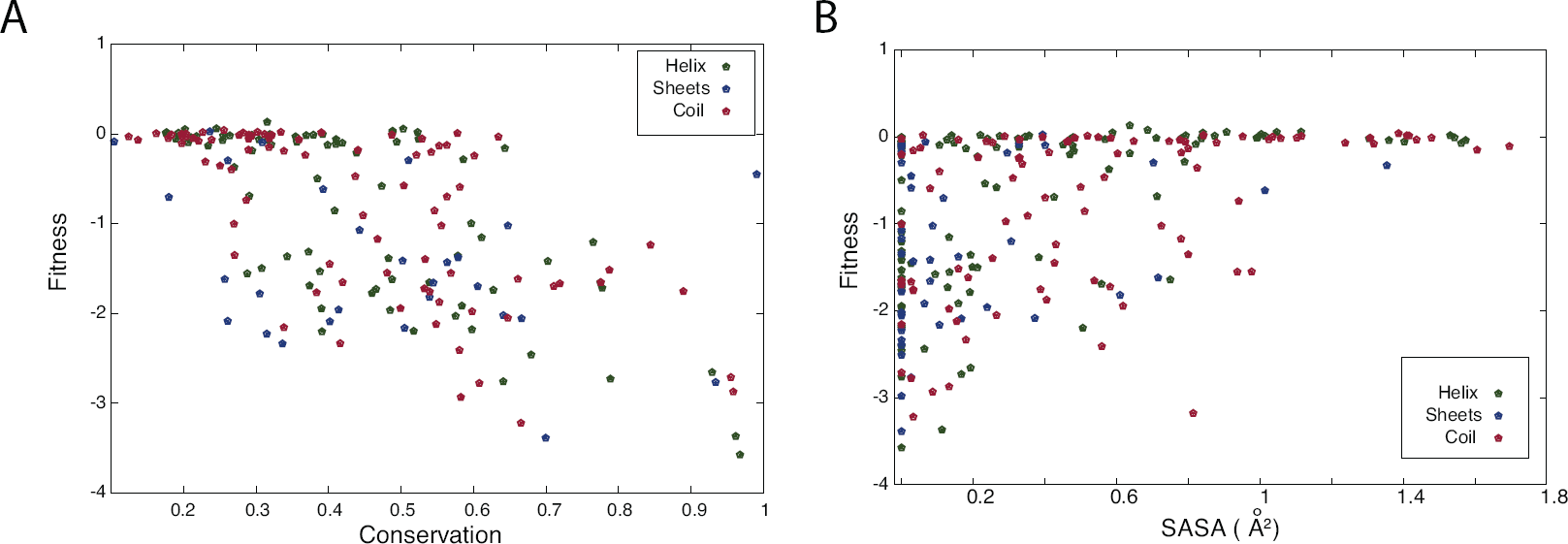
(A) Correlation between fitness and conservation (B) SASA and fitness shown only for alanine scan

### Dihedral angles

The dihedral angles *Φ* and *Ψ* suggest whether the conformation of amino acids are allowed or not. We examined if the dihedral angles predispose the amino acids to high or low sensitivity to mutations. However, the structure of the protein is available only for the wild type, and it is not also possible to easily guess if the mutations lead to a change or loss of structure. So, in this analysis, we use the structure and dihedrals corresponding to the wildtype as a reference. With these dihedrals, we represent the average and standard deviation of the relative fitness values for the 20 amino acids at the site in Supplementary Fig. 4. The figure suggests that the dihedrals in the wildtype do not predispose the amino acid to high or low fitness sensitivity.

**Supplementary Fig. 4:**
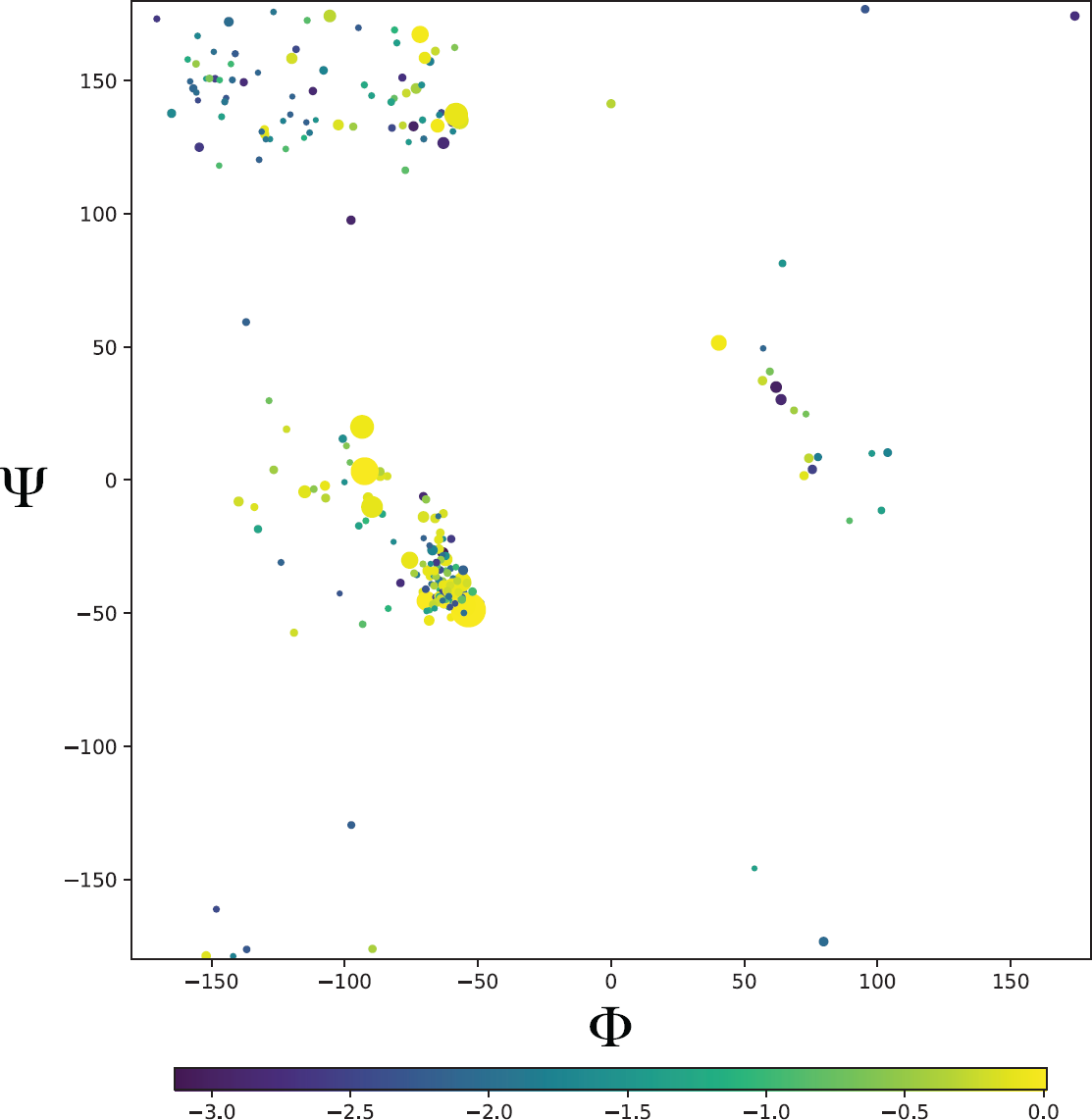
Dependence of fitness on Ramachandran angles. The color represents the average fitness change for the specific amino acid. Size of the data point is inversely proportional to the standard deviation of the fitness effects of 19 substitutions.

### Amino acid substitution matrix

The properties of wild type amino acid like solvent accessible surface area, conservation, and dihedrals examined so far are independent of the specific substitutions being made. These analyses can thus comment mainly on the average properties upon substitution, but miss the direct information about the substitution. To capture the specific mutation information, we use the position specific scoring matrix (PSSM) as a reference, which has been developed by combining the knowledge about all possible substitutions. PSSM score was calculated using PSI-BLAST for the multiple sequence alignment (MSA) The representation in Supplementary Fig. 5 shows a dependence, albeit weak.

**Supplementary Fig. 5:**
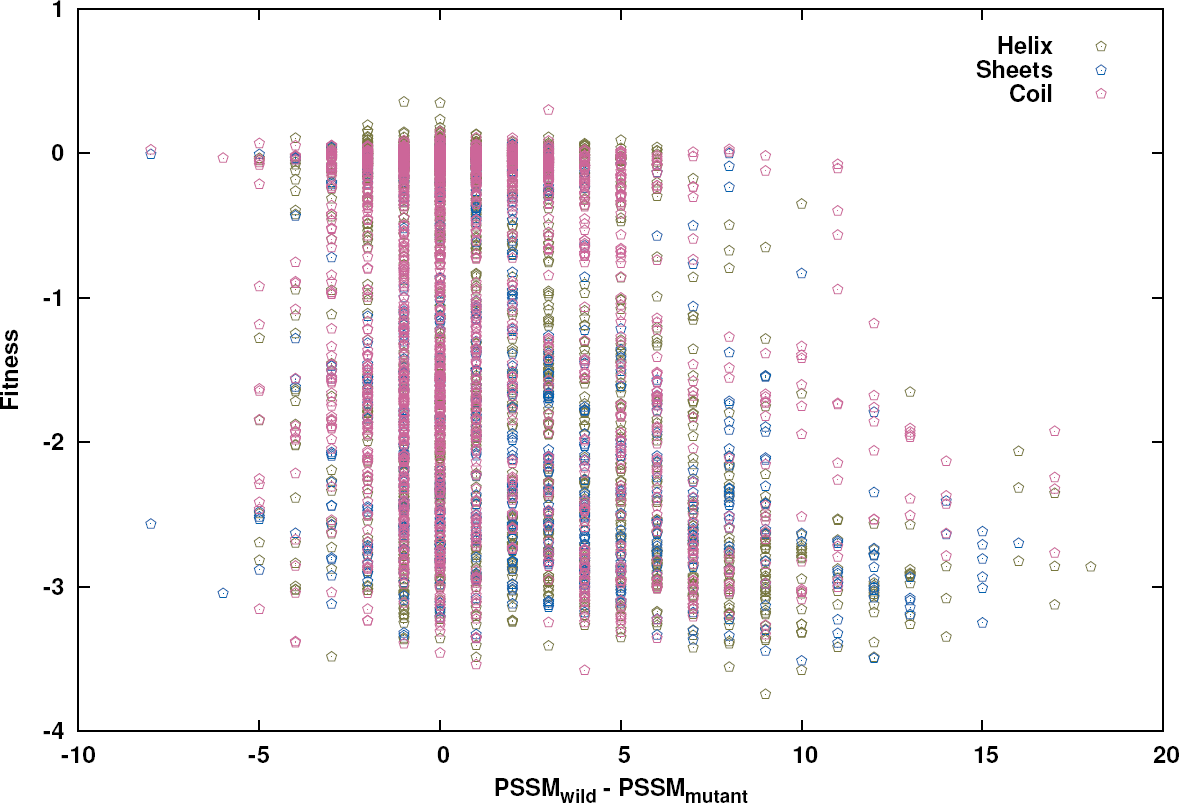
Comparison of fitness with Δ*PSSM* (= *PSSM_wildtype_ −PSSM_mutant_*) obtained from the PSSM. No strong pattern was observed.

### Spatial distribution of mutational sensitivity

In Fig. 4 of the manuscript, we explored how the comprehensive fitness data could be clustered to examine the patterns of sensitivity. The clustering simultaneously allowed the substitutions according to their charge type and other amino acids as well as by their residue numbers. Here we performed a simpler analysis which is much more intuitive. We filtered the comprehensive data for residues into two groups - which are extremely sensitive to all substitutions and which are not sensitive to any substitution. While the method of identification was different from the clustering, these amino acids in these two groups also appeared in the high and low sensitivity clusters.

**Supplementary Fig. 6:**
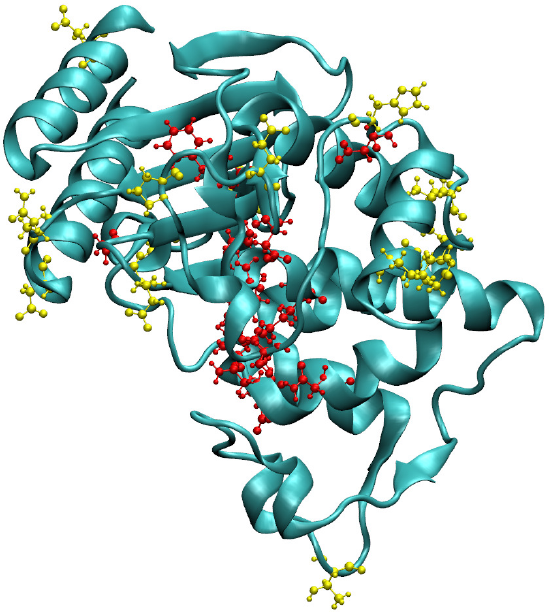
The most sensitive (all 19 substitutions lead to a fitness compromise *F* < −1.5 shown in red colour) and the least sensitive (all 19 substitutions are neutral(*F > −* 0.5) shown in yellow) positions are highlighted in the 3D structure of *β*-lactamase.

### Number of inter-residue contacts

Inter-amino acid interactions mediated by hydrogen bonds, salt bridges, stackings, etc determine how much a substitution disturbs the overall structural stability and function. While the biochemical details of the different interactions may be explored, and whether or not the nature of the substitutions conform with the existing interactions may also be investigated, a simpler metric is the total number of inter-residue interactions any given residue is involved in. We studied this by exploring atom level interactions with a 4 Å cutoff. The representation in Supplementary Fig. 7 shows that there is no relation.

**Supplementary Fig. 7:**
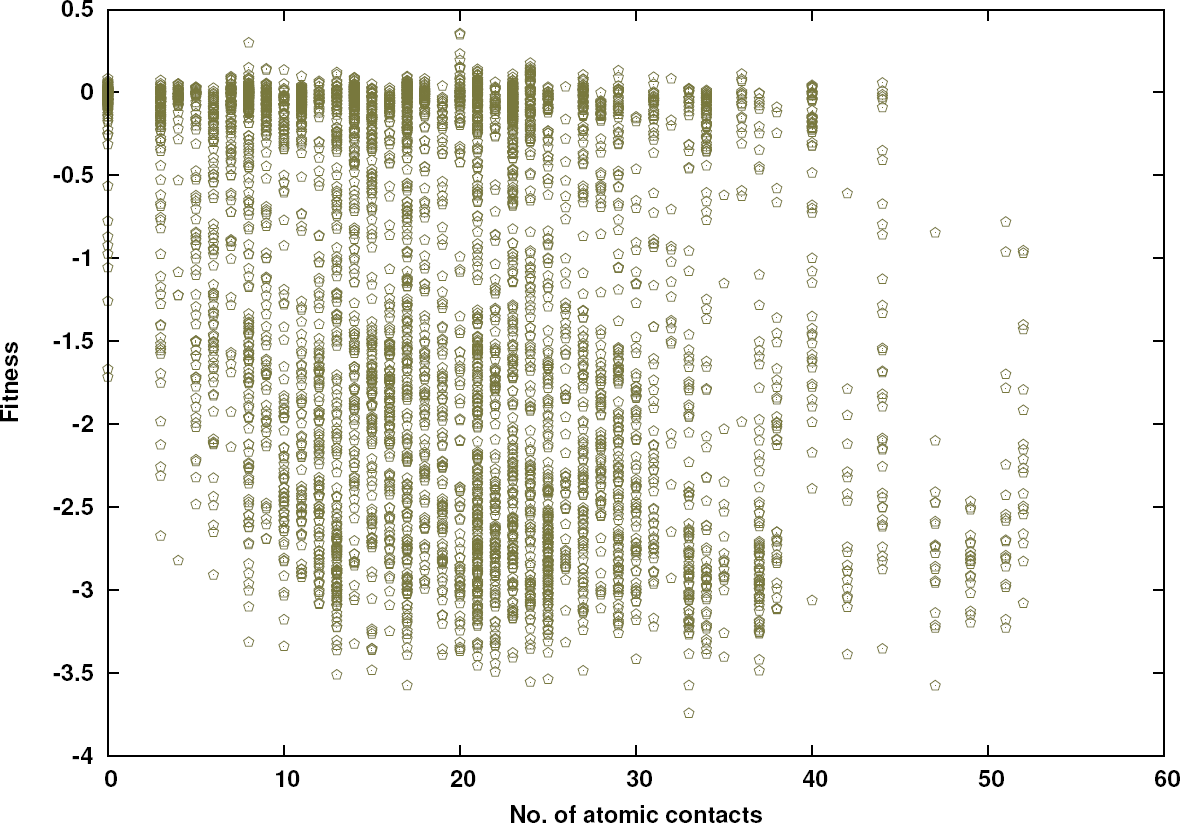
The effect of packing was studied by plotting the number of contacts relative to the fitness. No strong relation was observed. Fitness effects of substitutions of residues with large number of contacts are higher.

### Stability vs. Fitness

In order to explore the relation of sensitivity to substitutions and the thermodynamic stability an amino acid contributes to, we performed a simple thermodynamic analysis. The two underlying assumptions are that prevalence of a type of amino acid in the secondary structural elements reflects its contribution to the structural stability, and that the thermodynamic stability is suggestive of the fitness effects of the mutations. We used the data from the prevalence of amino acids in *α* helix ^27^ and *β* sheets ^26^ and plotted the ΔΔ*G* thus obtained with the relative fitness upon mutation. There was no correlation, and at this stage we do not know which of the two assumptions is not justified.

**Supplementary Fig. 8:**
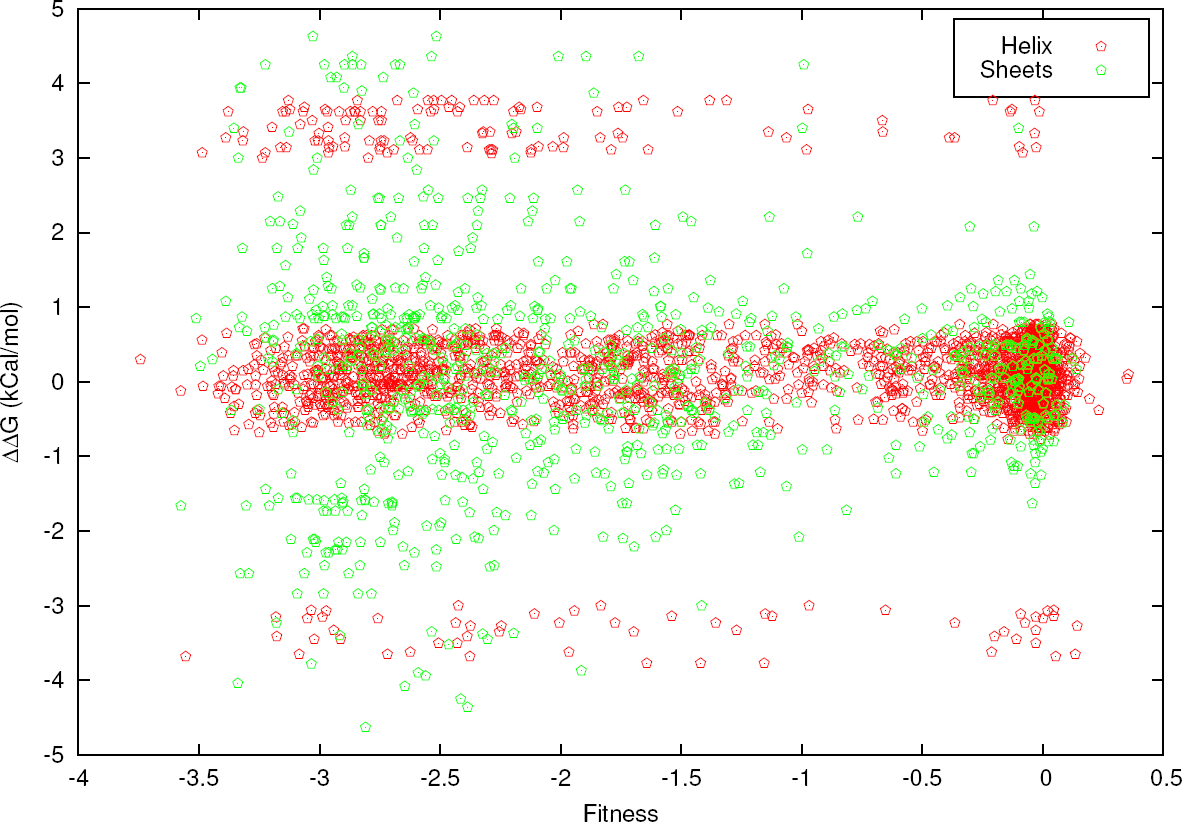
Role of stability. The change in stability, induced in the *α*-helix and *β*-sheet was calculated from the compiled data sets on the contributions of the amino acids to different secondary structural elements. No correlation with the expected change in stability and function were found.

### Salt bridges

Continuing with the emphasis on structural stability, we explored if the amino acids involved in salt bridges have a distinguishable sensitivity to mutations. In Supplementary Fig. 9 below, the analysis is presented pairwise for the different salt-bridges. It can be seen that the pattern is not common across all salt bridges.

**Supplementary Fig. 9:**
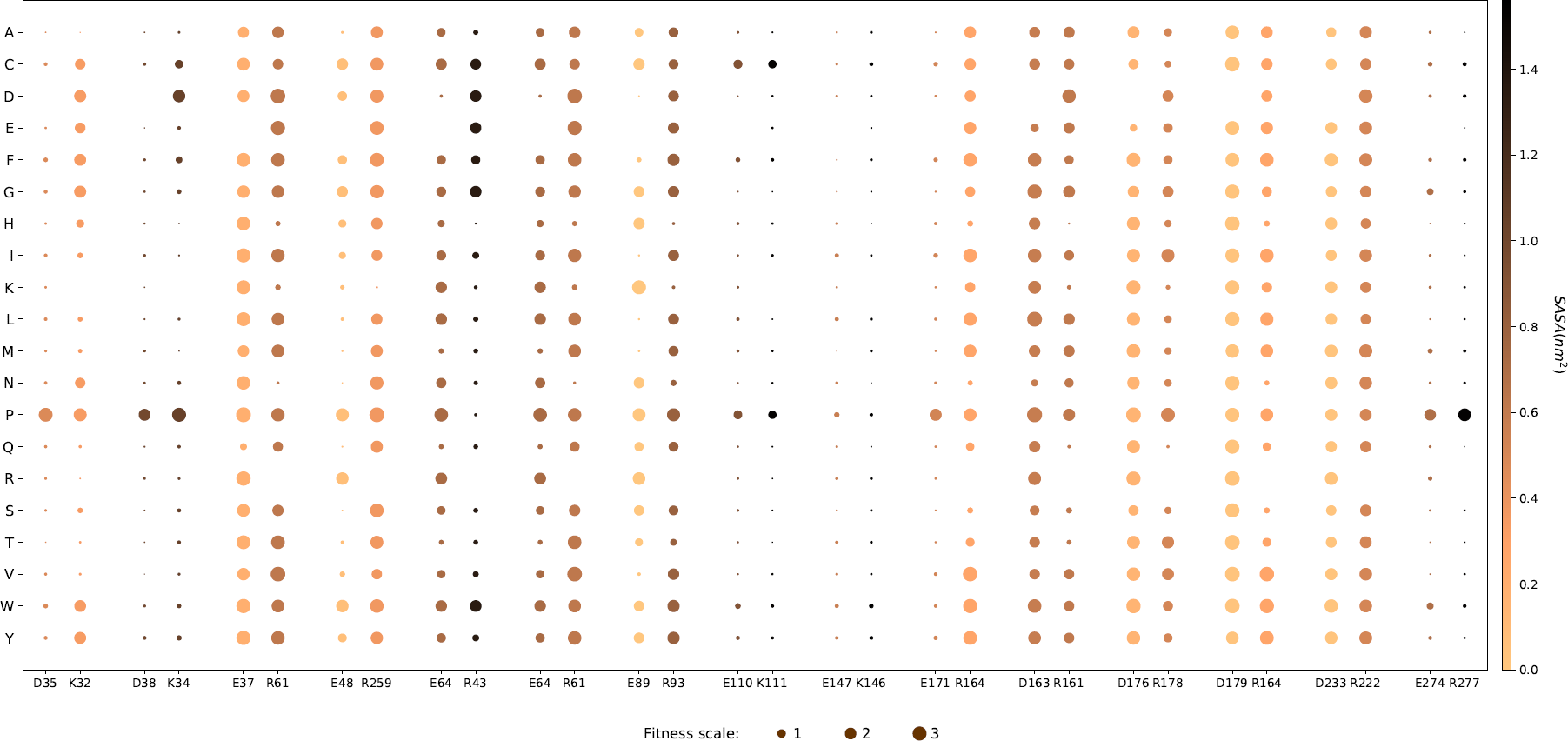
Fitness effects of salt bridge forming residues. The pairs of residues forming salt bridge are plotted close to each other. Size of the point shows the magnitude of fitness effect and increasing darkness of the points represents increasing SASA of the wild-type amino acid.

### Lysines: salt bridges vs. proximity to catalytic site

To weigh the differences between functional proximity and a comparison to structural stability via salt bridge formation, we compared lysines which are involved and not involved in salt bridges. The more notable patterns emerge from the lysines that are nearer to catalytic center, rather than from the differences in the involvement in salt bridges.

**Supplementary Fig. 10:**
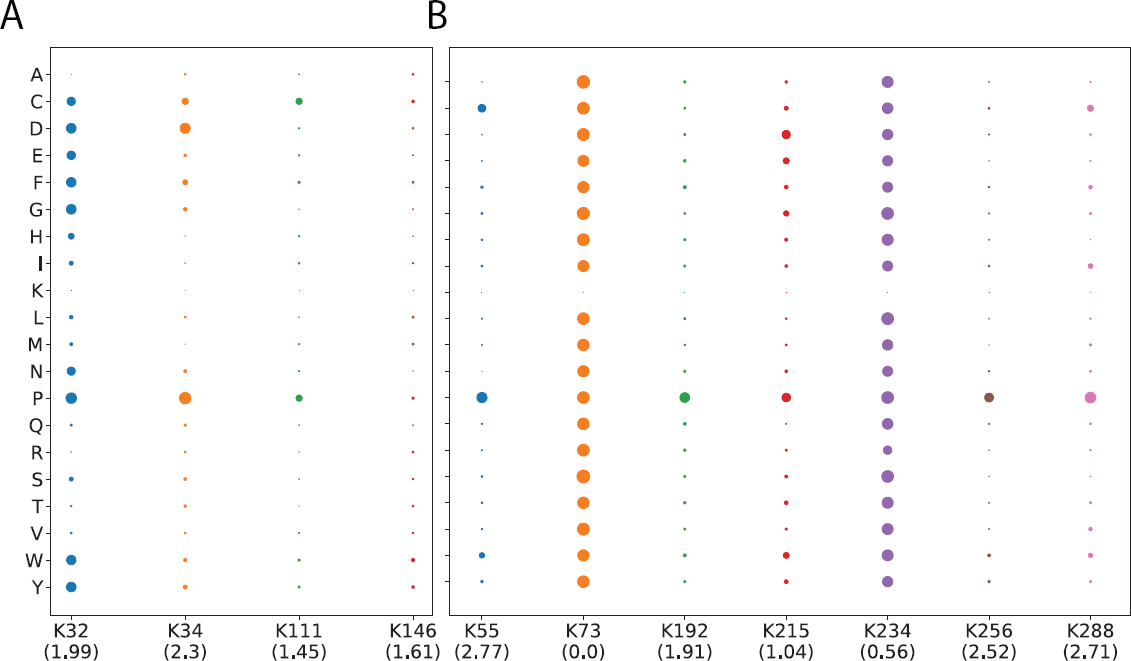
Comparison of functional importance of lysines which form salt-bridge with the ones that do not form. Comparison of functional importance of lysines which form salt-bridge with the ones that do not form: Effect of mutations of lysines (A) involved and (B) not involved in salt bridges. The number in the paranthesis indicates the distance in nm of the amino acid from the catalytic site. It is clear that sensitivity to mutations is higher for amino acids proximal to catalytic site than those forming salt bridges.

### Dose-response

The experimental data on the relative fitness with amino acid substitutions was characterized at 6 different drug concentrations and allowed for a comparison across the drug concentrations. Unlike the comprehensive fitness map, where the number of residues (about 263 of them) in one direction and the number of substitutions (19 of them) on the other direction, there is yet another dimension with the 6 external conditions. We converted this information again into a two dimensional map, with the 263 *×* 19 along one direction and the 6 drug concentrations along the other. The data was then clustered to identify groups of amino acid *subsitutions* which behaved similarly across all the external conditions. The groups that were identified with this clustering were represented in Fig. 7 of the manuscript.

**Supplementary Fig. 11:**
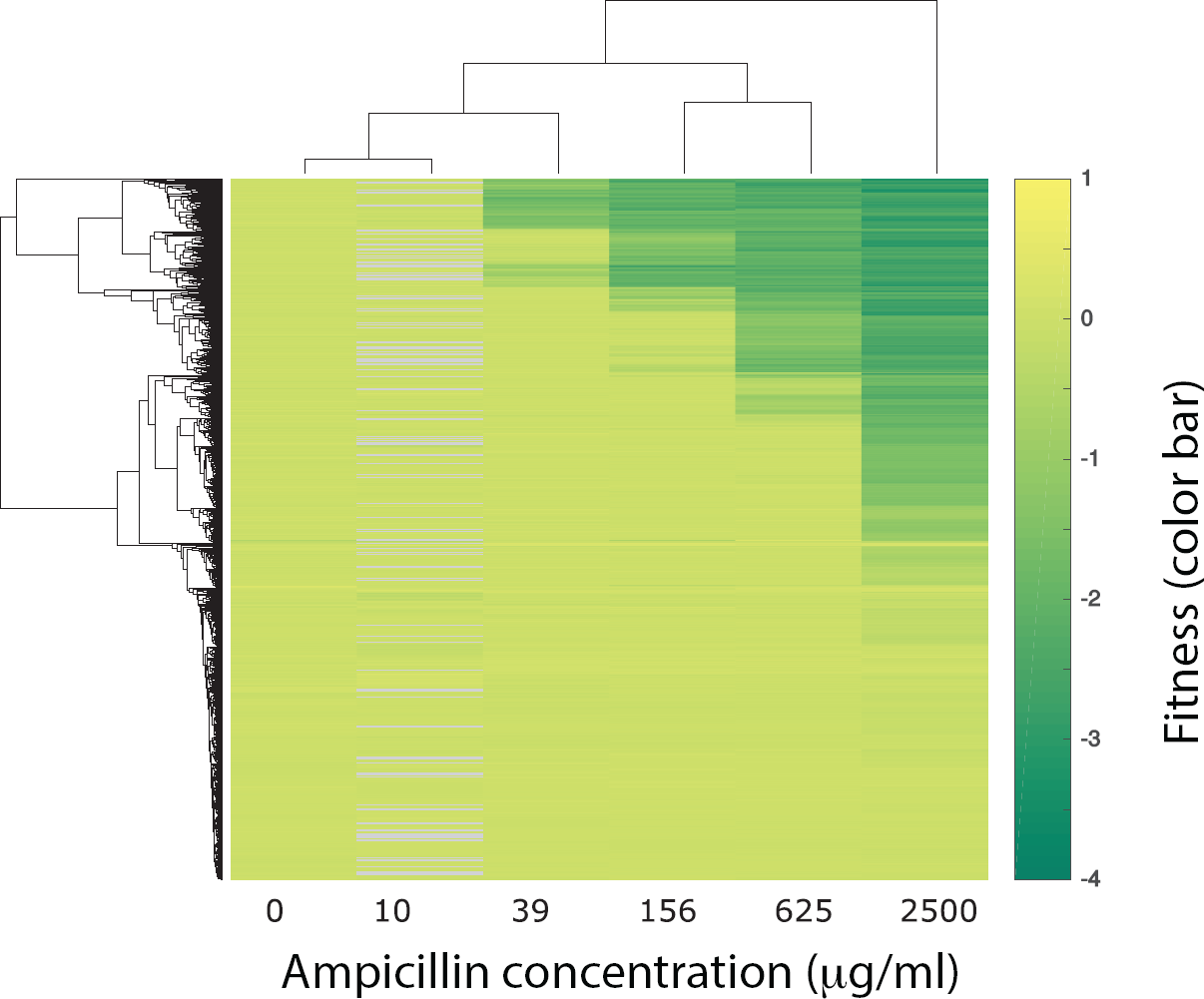
Clustering of all the point mutations based on the response to different concentrations of ampicillin

### SASA-Catalytic distance-BLOSUM score-Fitness

Two dimensional representations, with the variation in a parameter along one axis and its effect on the fitness are easier to represent and interpret. However, relative fitness simultaneously depends upon multiple factors, and a projection onto two dimensions will show patterns which may be confusing. So, we also tried to represent the data in a higher dimensional map for simultaneously analysing several dependences. However, we notice that the most significant dependence comes from the SASA.

**Supplementary Fig. 12:**
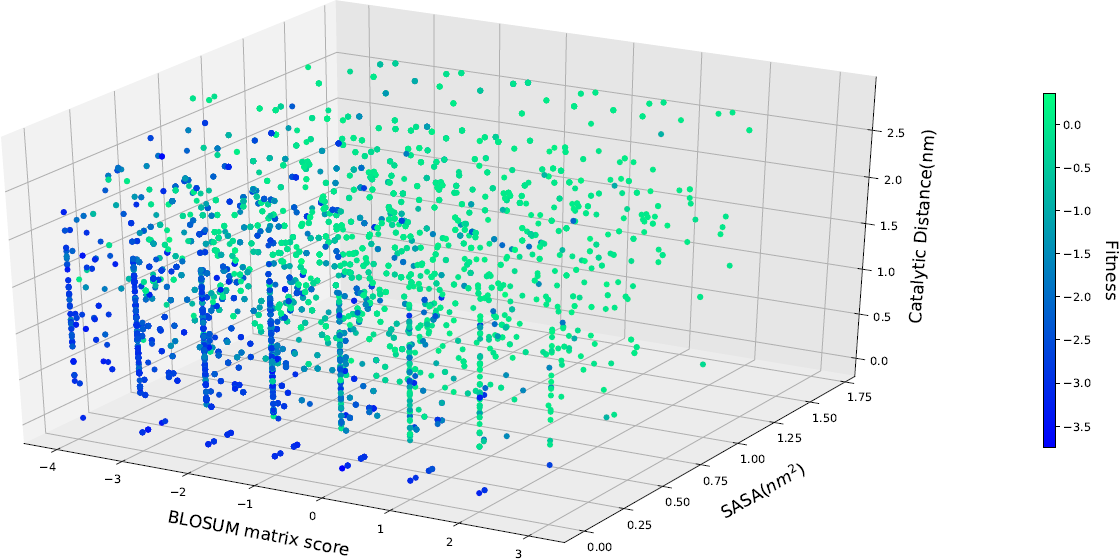
3D colormap showing the dependence of fitness on three variables simultaneously: BLOSUM matrix score, catalytic distance and SASA. Fitness is shown in color scale and the segregation of color indicates that these three variables together are able to classify the fitness effect as deleterious and neutral.

